## Supplementary tables and figure for "The Lipocone Superfamily: A Unifying Theme In Metabolism Of Lipids, Peptidoglycan And Exopolysaccharides, Inter-Organismal Conflicts And Immunity"

#### SUPPLEMENTARY MATERIAL

CONTENTS

Supplementary Table S1.
Supplementary Table S2.
Supplementary Figure Legends.
Supplementary Figures S1-S9.
Supplementary Data.
    List of identified genome contexts.
    yaml file of functional designations.

**Supplementary Table S1**. Lipocone family conserved contextual associations across distinct functional themes.

| Functional theme | Lipocone family | Genome associations[1] | Phyletic pattern notes[2] |
|---|---|---|---|
| Lipid head group exchange reactions | Euk-PTDSS1/2 | Phosphatidylserine production | Pan-eukaryotic |
| | Prok-PTDSS | Archaeophosphatidylserine production | sporadic archaea, bacteria, and viruses |
| | cpCone-1 | Kleisin-ScpA, HTH-ScpB | Patescibacteria |
| | | TM+LemA, TM+TM+MPTase, single cpCone-1 KptA domain fusion | |
| | | LP+LolA, 4TM with β-curl insert, [TM+NLPC/p60] | FCB group |
| | cpCONE-DUF2585 | HslV-Peptidase, GNAT, ClpB-HslU | Alphaproteobacteria |
| | | Glycolate-oxidase-GlcE, glycolate-oxidase-GlcF | |
| Cardiolipin synthesis | Wok-DUF2238 | Synaptojanin-like, [GlycosylTFase-A], [α/β-hydrolase], [NUDIX] | Terrabacteria, Pseudomonadota |
| | | Two-gene associations with diverse phosphatases: Calcineurin, HAD, PAP2 | Terrabacteria (sp), other lineages |
| Modified isoprenoid lipid synthesis | Wok-DUF2238 | Carotenoid biosynthesis module, GlycosylTFase-A, NUDIX | Actinomycetota |
| | YfiM-1 | amidophosphoribosyltransferase, UbiA-prenylTFase, RidA deaminase, TM-containing DUF5638, HD, PHP+Ig | Calditrichota, FCB group, Gemmatimonadetes |
| Lipid head group modifications in peptidoglycan dynamics | VanZ-1 & VanZ-2 | peptidoglycan glycosylTFases, D-Ala-D-Ala-peptidases, MurE-synthetase MurD-synthetase, MATE-flippase | Gene neighborhoods in Terrabacteria, FCB group, and Pseudomonadota |
| Lipid head group modifications in exopolysaccharide metabolism | VanZ-1 & VanZ-2 | WaaL-ligase, Wzc-CpsD-N, ElyC | Gene neighborhoods in Terrabacteria, FCB & PVC groups, and Pseudomonadota |
| Uncharacterized modifications of peptidoglycan and the outer membrane | VanZ-1+VanZ-i | Ferredoxin, α/β-hydrolase lipase, D-Ala-D-Ala-MβL transpeptidase | Betaproteobacteria |
| | VanZ-2 | SprA-N, GCV-H, 2TM+proline-rich-linker+TonB-C, TonB-C+OMP-β-barrel | FCB group |
| | VanZ-2 | ABC ATPase transporter, TM+coiled-coil+Papain-like or gly-gly-peptidase, SP+LTDs | Patescibacteria |
| Lipocone domains operating in or in transit to the outer membrane | YfiM-Griddle (up to 3 copies) | OMP-β-barrel(f), extended β-hairpin(f), LolA, POTRA, PLUG, TolB-N, Patatin lipase, GlycosylTFase-B, PAP2, LP+Synaptojanin, R-P(f) | Gram-negative bacteria |
| | YfiM-DUF2279 | OMP-β-barrel(f), MltG-endolytic-TGase, LP+Cytochrome-C7, PMM/PGM, GNAT, diaminopimelate-epimerase, Lysozyme | FCB group |
| | YfiM-DUF2279 | OMP-β-barrel(f), GlycosylTFase-A, OMP-β-barrel, SP+PDZ+ClpP-protease | FCB group |
| | ClaspCone-1 | TM(f) or 5TM(f), TULIP(f) or Ig and MPTase(f), [PHP](f), GDSL-Lipase, MBOAT | Pseudomonadota, Planctomycetota |

| | | | |
|---|---|---|---|
| Membrane-anchoring linkage | Skillet-1 | Specialized lipobox(f), diverse ligand-binding domains(f): Ig, Jellyroll, βPs, Concanavalin, OB-fold, SHOCT, MORNs | Bacillota, FCB group, Pseudomonadota |
| Lipid-associated signaling systems, standalone proteins | VanZ-1 | HTH(f), RHH(f), YycI(f), RDD(f), Glyoxylase(f), NPCBM(f) | Widespread, sporadic linkages |
| | VanZ-2 | cNMPDB(f), FHA(f), KTSC(f), Papain(f), TPRs(f), Calcineurin(f), CBD9(f) | |
| | Skillet-3 | Ig(f), β-sandwich(f), helix-grip(f), βPs(f), MORNs(f), Lipocalin(f), β-barrel(f) | Pseudomonadota, FCB group, Terrabacteria |
| Lipid-associating signaling systems, multicomponent | VanZ-1 | HAAS(f), PadR-HTH | Bacillota |
| | Skillet-2 | helix+TM or ZnR+helix+TM or HTH+L12-ClpS+TM, TetR transcriptional repressor, [HMG-CoA-reductase+GHMP-kinase], [SP+Ig repeats] | Bacteriodota, Bacillota (sp) |
| | Skillet-DUF2809 | wHTH, cytoplasmic-helix+6TM protein. Joined by one or more of: ElyC, CreD, Coq4, Lcp-like, DUF1361, TGase | FCB group, Pseudomonadota (sp) |
| Antiviral immunity | Min-Wnt | DUF3892(f) | Pseudomonadota (sp) |
| | | 3-strand β-meander(f), LP+PPTs, SP+Glycosyl-hydrolase, SP+β-helix | Bacteroidota |
| | | Standalone | Cyanobacteria |
| | | helical-domain+PcfJ-GNAT(f) | Duplodnaviria |
| Toxin domains in polymorphic and allied conflict systems | Min-Wnt | SP or LP+tail(f), LP+Imm-BamE or LP+Imm-Jellyroll or Imm-4TM | Terrabacteria, Pseudomonadota, FCB group, Elusimicrobia, Acidobacteria, PVC group (sp), Archaea (sp) |
| | | Polymorphic toxin delivery systems: T1SS, T4SS, T6SS, T7SS, T9SS, DUF4157-MPTase, Immunity proteins as above | |
| | | LP+Cystatin-FD (f), LP+Imm-Jellyroll (dominant) or LP+Imm-BamE | Bacteroidota |
| | Prok-SAA | Polymorphic toxin delivery systems: T6SS, MuF, TM+[TM+TM+]Imm-SAA or LP+Imm-BamE | Spirochaetota, Nitrospirota, Acidobacteriota, Terrabacteria (sp), PVC group, Pseudomonadota (sp), Fusobacteriota, Bacteriodota (sp) |
| | | PGBD(f), TM+Imm-SAA | Pseudomonadota (sp) |
| | Prok-TelC | Polymorphic toxin delivery systems: T6SS, T7SS, MPTase-DUF4157, ZU5+vWA core {31064832}, Imm-TipC, Imm-Zu5/vWA | Bacillota, Actinomycetota (sp), Myxococcota (sp), FCB and PVC groups (sp), Pseudomonadota (sp) |
| | | SP+GbpC+MucBP-IG(f), Imm-TipC | Bacillota and Actinomycetota (sp) |
| | | TPM+TPM(f) or TPM+Ig, Imm-4TM | Bacteriodota |
| | CapCone-1 | Polymorphic toxin delivery systems: T6SS (including PsbP/MOG1-like fusion), MPTase-DUF4157, LP+Imm-BamE | Pseudomonadota, PVC group, Terrabacteria (sp), FCB group (sp) |

| | | | |
|---|---|---|---|
| | | Cystatin-FD+linker, LP+Imm-BamE | Bacteroidota |
| | CapCone-2 | SP(f), SP+Imm-BamE | Bdellovibrionota, Acidobacteria (sp) |
| | | Polymorphic toxin delivery systems: T6SS, MPTase-DUF4157 | Pseudomonadota (sp), FCB group (sp), PVC group (sp), Archaea (sp) |
| | | ANKs(f), SP+Imm-SAS6-N-like-β-sandwich | PVC group (sp), Pseudomonadota (sp) |
| | ClaspCone-2 | Polymorphic toxin delivery systems: T6SS, Imm-4TM | FCB group (sp), PVC group (sp), Pseudomonadota (sp) |
| | VanZ-1 | Polymorphic toxin delivery systems: T6SS | Bacillota |
| Toxins in predator-prey and other inter-specific conflicts | Min-Wnt | SP+half-β-barrel(f), CC-motif-containing-tail(f), C-terminal helical-extension(f) | Bacteroidota, PVC group, Terrabacteria (sp), Hemichordata, Rotifera, fungi |
| | | Broken-hairpin(f) | Alphaproteobacteria (sp), Duplodnaviria (sp), Terrabacteria (sp) |
| | CapCone-2 | SP(f), Patatin(f), Lipocalin, acyltransferase+TM+TM+TM, SP+α/β-hydrolase, SP+OMP-β-barrel | Bdellovibrionota, Holophagales, Archangium, Woeseiaceae, Labrenzia, Roseibium |
| | Prok-SAA | SP+MTPase+Prok-SAA+vWD+Ig+Ig | Gemmatimonadetes (sp), Pseudomonata (sp) |
| | Skillet | Histidine kinase-Receiver, MPTase, Papain-like, MTases, LysM & other ligand-bindings domains, etc. | Omnitrophica Patescibacteria |
| | Prok-TelC | NAGPA(f), ligand-binding(f): Ig, CW-repeats, βPs, β-sandwich, etc. | Bacillota |
| | | PGBD+PGBD+Rv2525c-like-TIM-barrel(f), SP+Ig+Ig*, 3TM-CCDN*, SP+VanY* | Bacillota, fungi (sp), Actinomycetota* (sp) |
| Resistance to antimicrobial agents | VanZ-1, VanZ-2, Skillet-DUF2809* | VanY, vancomycin resistance modules, D-Ala-D-Ala-MβL* | Terrabacteria, FCB group (sp)*, Pseudomonata (sp)* |
| | YfiM-1 | Thioredoxin, DTW-SPOUT, acetate—CoA-ligase+ATP-grasp+GNAT, HKD fold phosphatidylserine synthetase | Gammaproteobacteria (sp) |

[1](f): denotes a domain that is directly fused to the Lipocone family; *: associations present in a phylogenetically restricted subset; [x]: association is not universally observed; GlycosylTFase-A: glycosyltransferase-A; TFase: transferase; TGase: transglycosylase; CW: cell wall; MTase: Methylase; MBL: metallo-β-lactamase; TM: transmembrane; SP: signal peptide; LP: membrane-anchored lipoprotein; T[x]SS: Type-X-secretory system; Imm: immunity protein; βPs: β-propellers;

[2](sp): denotes sporadic distribution in the listed phylogeny; *: phylogenies with restricted associations

Supplementary Table S2. Significant enrichment of Lipocone family contextual associations across functional categories.

| Lipocone family | Function | p-value |
|---|---|---|
| Prok-TelC | Biological conflict | 0.0 |
| Min-Wnt | Biological conflict | 0.0 |
| VanZ-2 | Sugar metabolism | 0.0 |
| Skillet-1 | Adhesion/extracellular matrix | 0.000001 |
| VanZ-2 | Peptidoglycan | 0.000002 |
| YfiM-Griddle | Outer membrane | 0.000015 |
| VanZ-1 | Sugar metabolism | 0.000022 |
| YfiM-1 | Isoprenoid metabolism | 0.000171 |
| Wok-DUF2238 | Isoprenoid metabolism | 0.000242 |
| Skillet-DUF2809 | Exopolysaccharide | 0.000296 |
| Skillet-2 | Transcription | 0.00053 |
| CapCone-1 | Biological conflict | 0.000636 |
| VanZ-1 | Exopolysaccharide | 0.000836 |
| Skillet-3 | Adhesion/extracellular matrix | 0.000839 |
| cpCone-i | Cell membrane | 0.003064 |
| CapCone-2 | Biological conflict | 0.003375 |
| VanZ-i | Sugar metabolism | 0.003824 |
| cpCone-1 | Cell membrane | 0.007168 |

**Supplementary Figure Legends.**

**Figure S1**. Phyletic distribution patterns of Lipocone superfamily. (top) Percentage of genomes containing at least one representative of a given Lipocone family within a discrete phylogeny are reported as colored in the provided legend. (bottom) Bar graph depicting phyletic depth (bar height in $D_i$ - see methods) and breadth (bar width) for Lipocone clades (see Figure 4). Coloring contrasts membrane-associating clades with diffusible clades.

**Figure S2**. Structural representatives of the Lipocone superfamily. Representative structures or predicted models from Lipocone families. Core helices (H1, H2, H3, and H4) are colored uniformly across the structures as in Figure 1A. Loops and inserts are outlined and transparent, distinctive features are labeled as appropriate. Protein Data Bank (PDB) IDs or protein sequence identifiers used to generate AF models are provided. The core three ancestral active site positions (see Figure 2) are rendered as ball-and-stick, with carbons colored green and other atoms colored as standard element colors.

**Figure S3.** Critical difference diagram depicting group-wise differences across TM tendency score distributions in Figure 1C (see Methods). Groups connected by horizontal bars are not significantly different (Bonferroni-adjusted $p > 0.05$); groups not connected are significantly different (adjusted $p < 0.05$). Three non-linked groups are observed in this diagram: 1) those with clear negative TM propensity scores (containing families predicted to be soluble), 2) those with clear positive TM propensity scores (containing families predicted to insert into the membrane), and 3) those with borderline scores. The third category includes families known to insert into the ER membrane (PTDSS1/2) and those predicted to insert into the outer bacterial membrane (YfiM clade).

**Figure S4**. Structural diversity in the cpCone clade. Each panel depicts a distinct structural configuration observed in the cpCone clade, colored by rainbow palette from N- to C-termini (left structure) and by equivalent helix (right structure). All representative structures were selected from the cpCone-1 family, except where noted. AF models are based on sequence provided as NCBI accession number labels.

**Figures S5-S7.** Lipocone domain-centered subgraphs of contextual network in Figure 6. These subgraphs capture significant enrichment of different functional categories (Table S2). Subgraph network nodes, edges, scaling, and coloring as described in Figure 6 legend.

**Figure S8**. Multiple sequence alignment of serine-containing lipobox (SLP). Sequences are labeled to the left by NCBI accession number and organism abbreviations. The conserved serine residue position, denoted at the top of the alignment by an asterisk, is shaded in red, with text colored in white. Other residue positions are colored according to consensus conserved biochemical properties: hydrophobic (h) and aromatic (a) residues are shaded yellow, polar (p) residues are shaded blue, small (s) and tiny (u) residues are shaded green, and positively-charged (+) residues are shaded red. Diversity of domains Cterminally fused to the SLP are depicted to the right of the alignment, represented as geometric shapes. Organism abbreviations as follows: Obac: *Oscillospiraceae bacterium*; Cbac: *Clostridia bacterium*; Lbac: *Lachnospiraceae bacterium*; CEqu: *Candidatus Equihabitans*; Rzha: *Roseburia zhanii*; Rusp: *Ruminococcus* sp; Aaut: *Aceticella autotrophica*; Nthe: *Natranaerobius thermophilus*; CFim: *Candidatus Fimenecus*; Busp: *Butyrivibrio* sp; Eusp: *Eubacterium* sp; Rbac: *Ruminococcaceae bacterium*; Chsp: *Chryseobacterium* sp; Flsp: *Fluviicola* sp; Zpro: *Zunongwangia profunda*; Ibac: *Ignavibacteria bacterium*; Mbac: *Myxococcales bacterium*; Pbac1: *Planctomycetota bacterium*; Byua: *Bradyrhizobium yuanmingense*; Rhsp: *Rhodopseudomonas* sp; Afer: *Acidimicrobium ferrooxidans*; Bbac1: *Bacillota bacterium*; Etay: *Eisenbergiella tayi*; Rosp: *Roseburia* sp; Bbac2: *Betaproteobacteria bacterium*; Pbac2: *Pseudomonadota bacterium*; Bbac3: *Burkholderiales bacterium*; Idsp: *Ideonella* sp; Pisp: *Piscinibacter* sp; Aant: *Algoriphagus antarcticus*; Fbac: *Flavobacteriales bacterium*; Friv: *Flavobacterium rivulicola*; Mlut: *Mongoliitalea lutea*; Ga: *Gammaproteobacteria* ; Sysp: *Syntrophorhabdus* sp; Aadv: *Apibacter adventoris*; Bbac4: *Bacteroidota bacterium*; Chsp: *Chryseobacterium* sp; Spsy: *Sphingobacterium psychroaquaticum*; mbac: marine bacterium; Ster: *Sebaldella termitidis*; Abac: *Armatimonadota bacterium*; Bcla: BD1-7 clade; Gpen: *Gallaecimonas pentaromativorans*; Kgeo: *Kangiella geojedonensis*; Ktai: *Kangiella taiwanensis*; Posp: *Porphyromonas* sp; Xbre: *Xylanibacter brevis*; Pmul: *Prevotella multiformis*; Scop: *Segatella copri*; Bbac5: *Bacteroidales bacterium*; Masp: *Mariniphaga* sp; Pbac3: *Prolixibacteraceae bacterium*; Dbac: *Desulfobacteraceae bacterium*.

**Figure S9**. Structural overview of newly identified immunity proteins pairing with toxin-containing proteins in polymorphic and allied toxin systems. Top panel depicts concordance of core secondary structure elements across BamE-like immunity protein families, with N-terminal α-helix dyad colored in blue and green and the four strands of the core β-meander colored in a yellow, light green, orange, and purple order. Bottom panel depicts rainbow palette coloring of the Jellyroll domain-containing immunity protein and the 4-TM protein. The Immunity-SAA protein is colored by secondary structure element, with conserved cysteine residues rendered as ball-and-stick. Protein DataBank ID (PDBID) or AF model-generating sequence is provided.

**Figure S10**. Sequence and structure overview of the broken-hairpin domain. (A) Multiple sequence alignment of broken-hairpin domain, with conserved axR (with 'a' representing an aromatic residue, 'x' representing any residue, and R representing an arginine residue) motif positions labeled above alignment. Coloring and conserved consensus abbreviations as in Figure S4. (B) AF models of broken hairpin domain, loops colored in gray and strands in orange. axR motifs are rendered as ball-and-stick representations. (C) Selection of domain architectures observed with the broken-hairpin domain, arranged and labeled by general functional theme. Depictions and abbreviations as described in Figures 5 and 6 legends. (D-E) AF models exploring the positioning of the broken-hairpin domain relative to distinct N- or C-terminal effector domains.

### SUPPLEMENTARY FIGURE S1

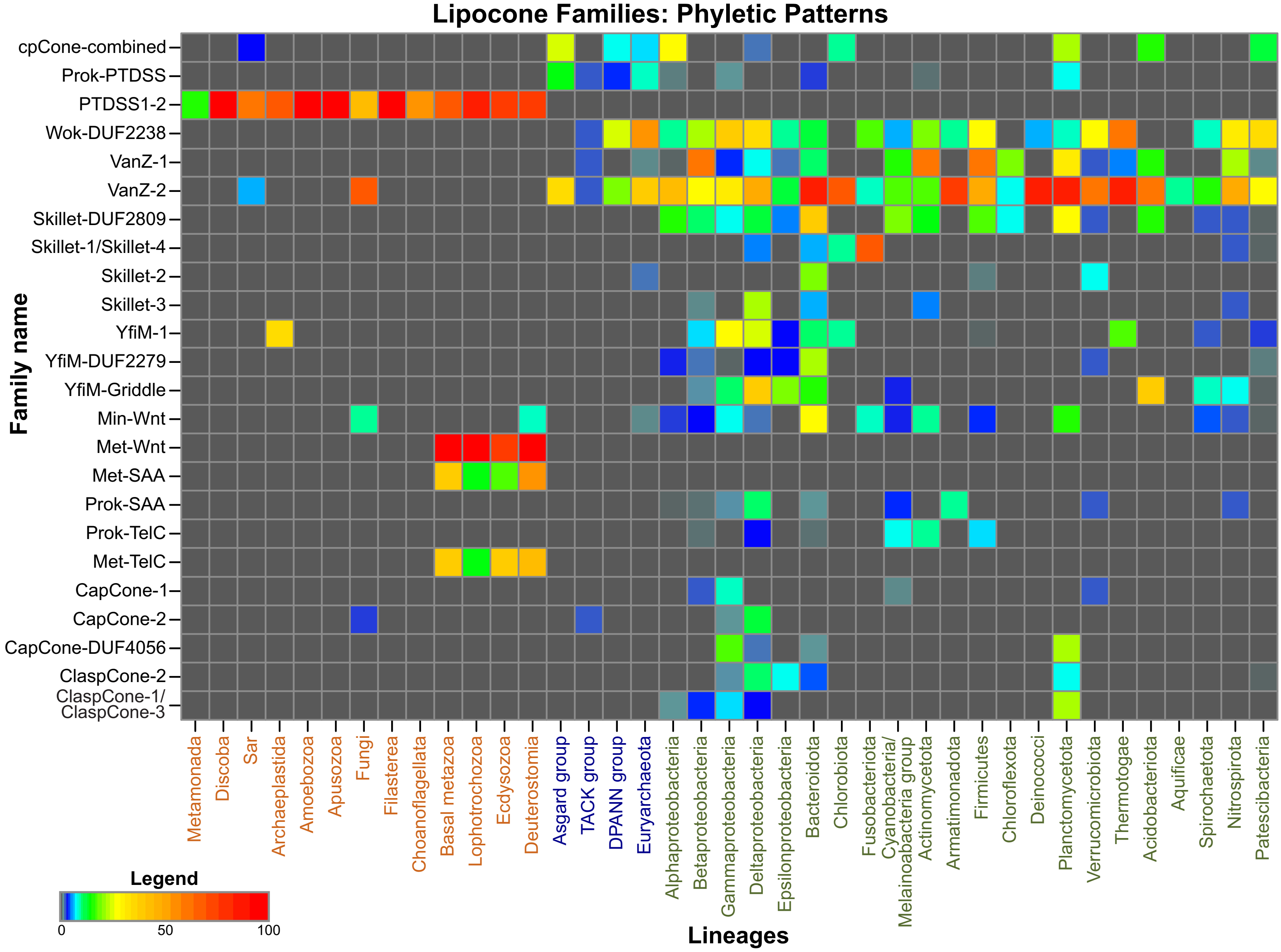


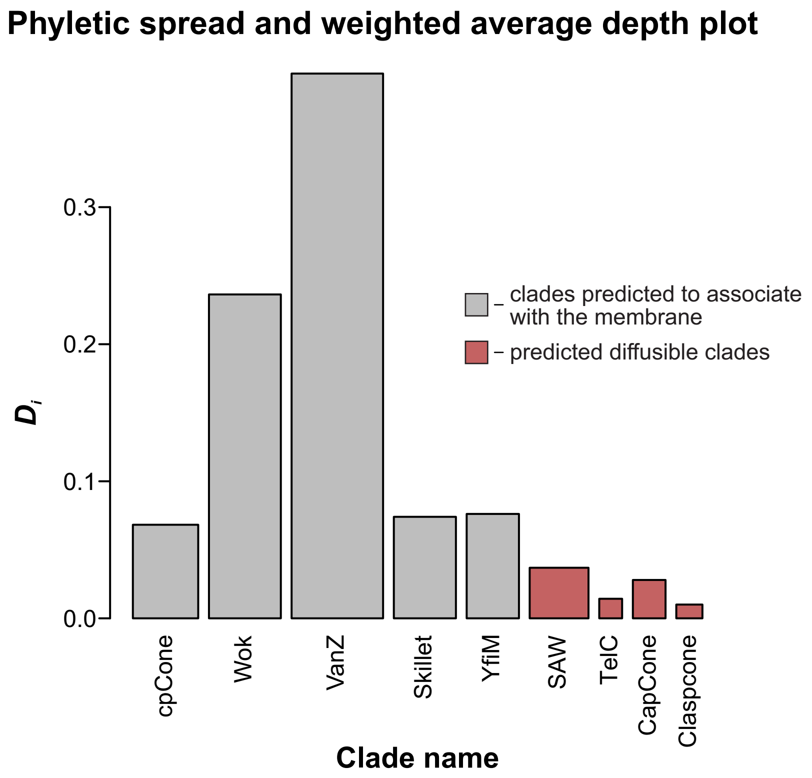

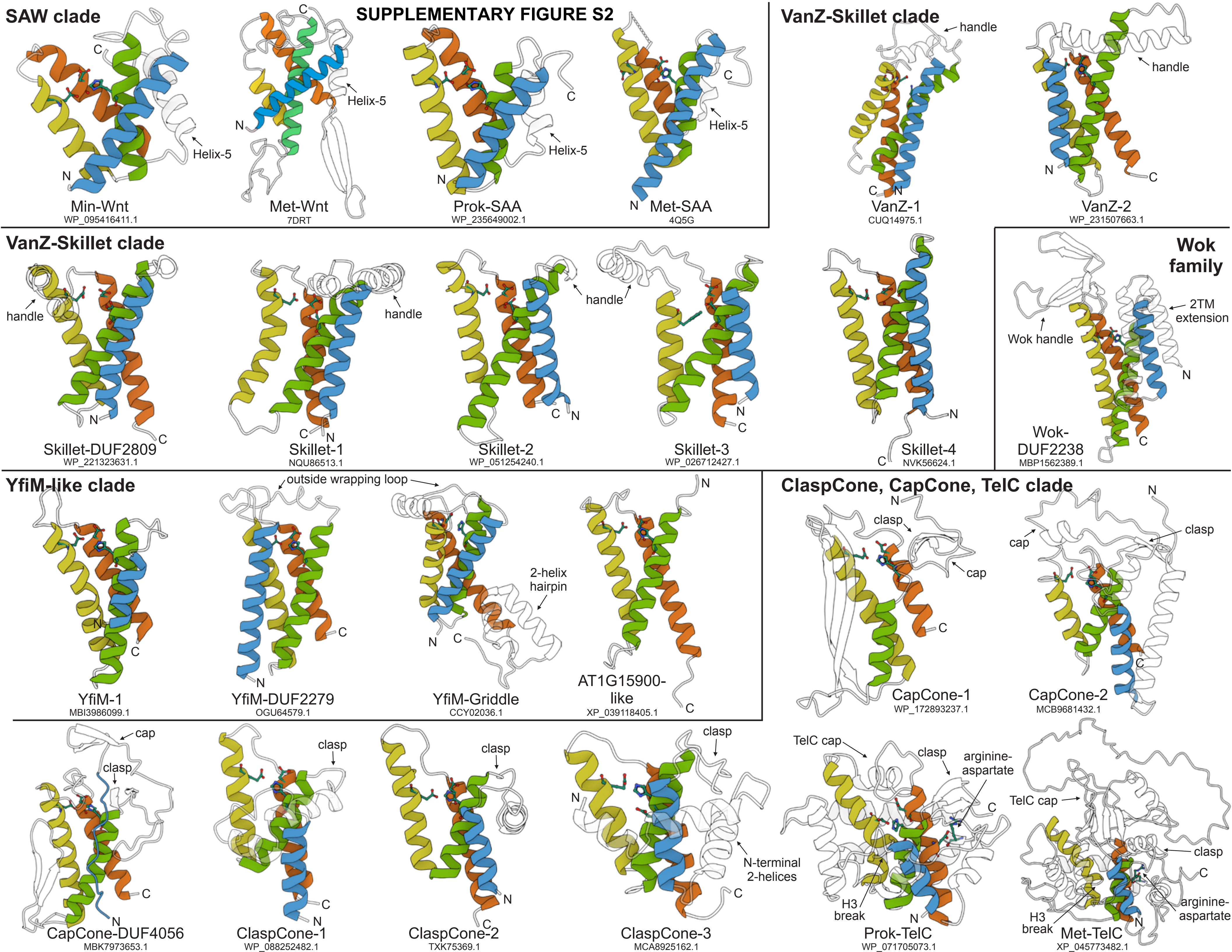
SUPPLEMENTARY FIGURE S2
SAW clade
Helix-5
Min-Wnt
WP_095416411.1
Met-Wnt
7DRT
Prok-SAA
WP_235649002.1
Met-SAA
4Q5G
VanZ-Skillet clade
handle
VanZ-1
CUQ14975.1
VanZ-2
WP_231507663.1
VanZ-Skillet clade
Skillet-DUF2809
WP_221323631.1
Skillet-1
NQU86513.1
Skillet-2
WP_051254240.1
Skillet-3
WP_026712427.1
Skillet-4
NVK56624.1
Wok family
Wok handle
2TM extension
Wok-DUF2238
MBP1562389.1
YfiM-like clade
outside wrapping loop
2-helix hairpin
YfiM-1
MBI3986099.1
YfiM-DUF2279
OGU64579.1
YfiM-Griddle
CCY02036.1
AT1G15900-like
XP_039118405.1
ClaspCone, CapCone, TelC clade
clasp
cap
CapCone-1
WP_172893237.1
CapCone-2
MCB9681432.1
CapCone-DUF4056
MBK7973653.1
ClaspCone-1
WP_088252482.1
ClaspCone-2
TXK75369.1
N-terminal 2-helices
ClaspCone-3
MCA8925162.1
TelC cap
arginine-aspartate
H3 break
Prok-TelC
WP_071705073.1
Met-TelC
XP_045773482.1

**SUPPLEMENTARY FIGURE S3**

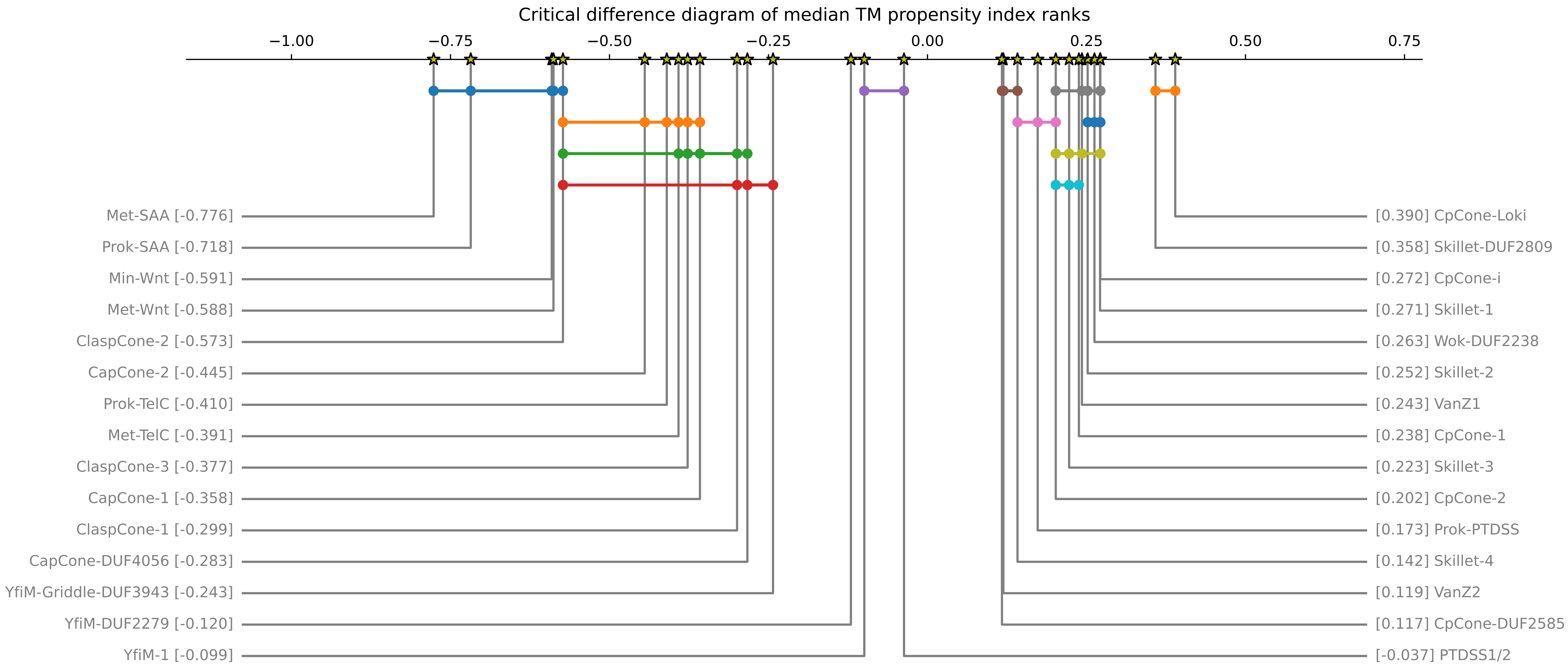

### S4. Structural diversity in the cpCone clade

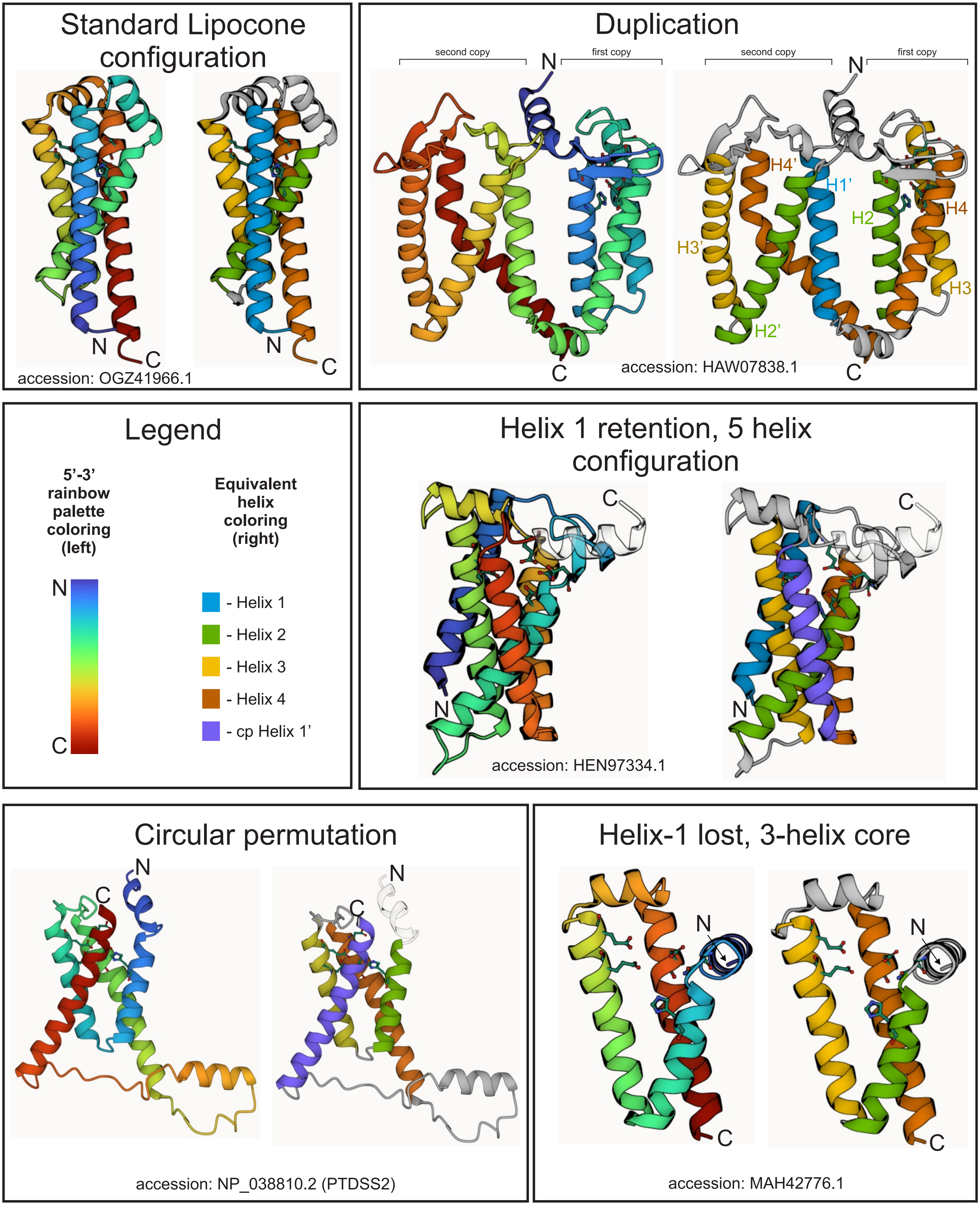

### SUPPLEMENTARY FIGURE S5.

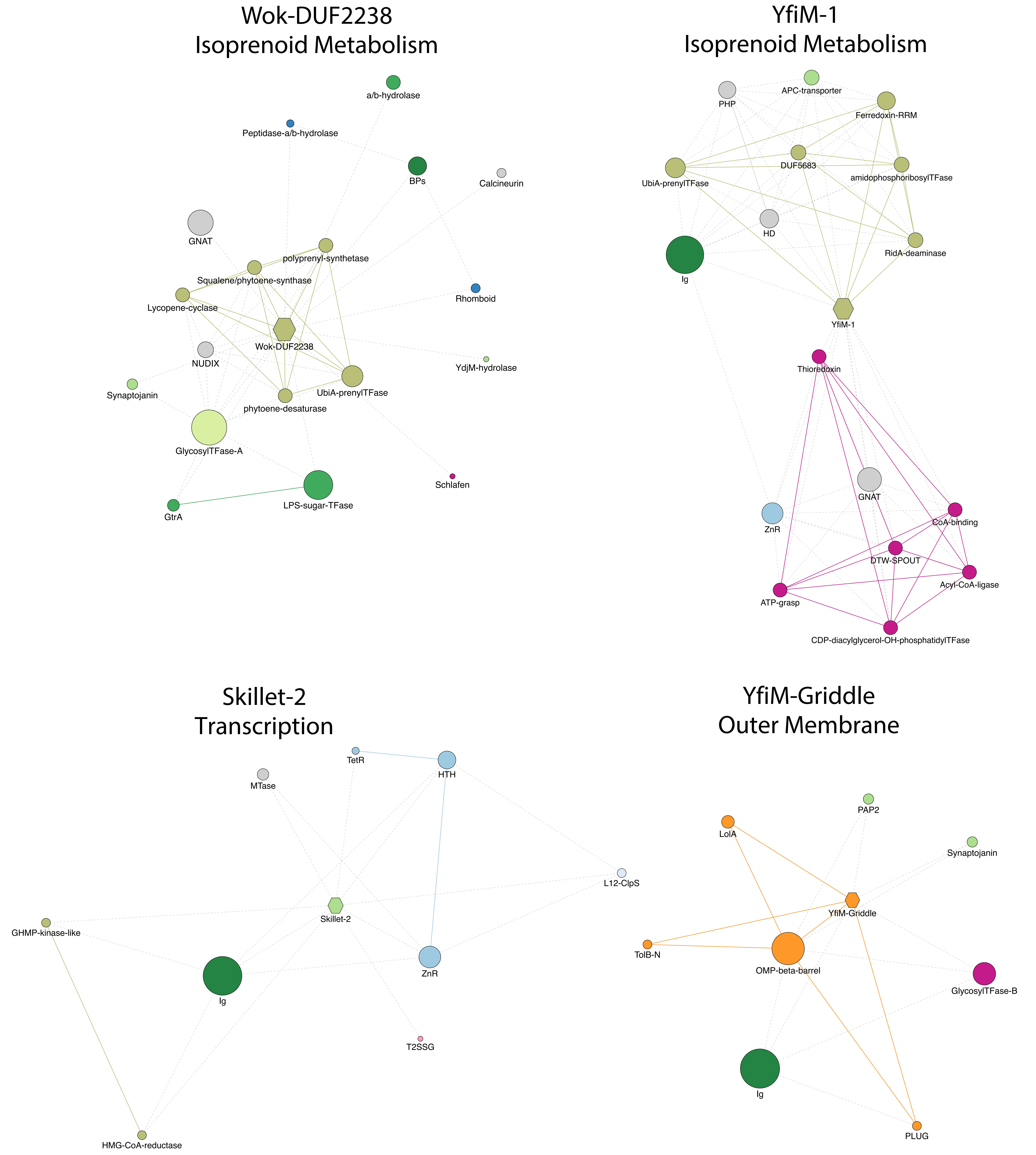

#### SUPPLEMENTARY FIGURE S6.

Sugar metabolism, Adhesion/extracellular matrix, Peptidoglycan, and Exopolysaccharide subgraphs

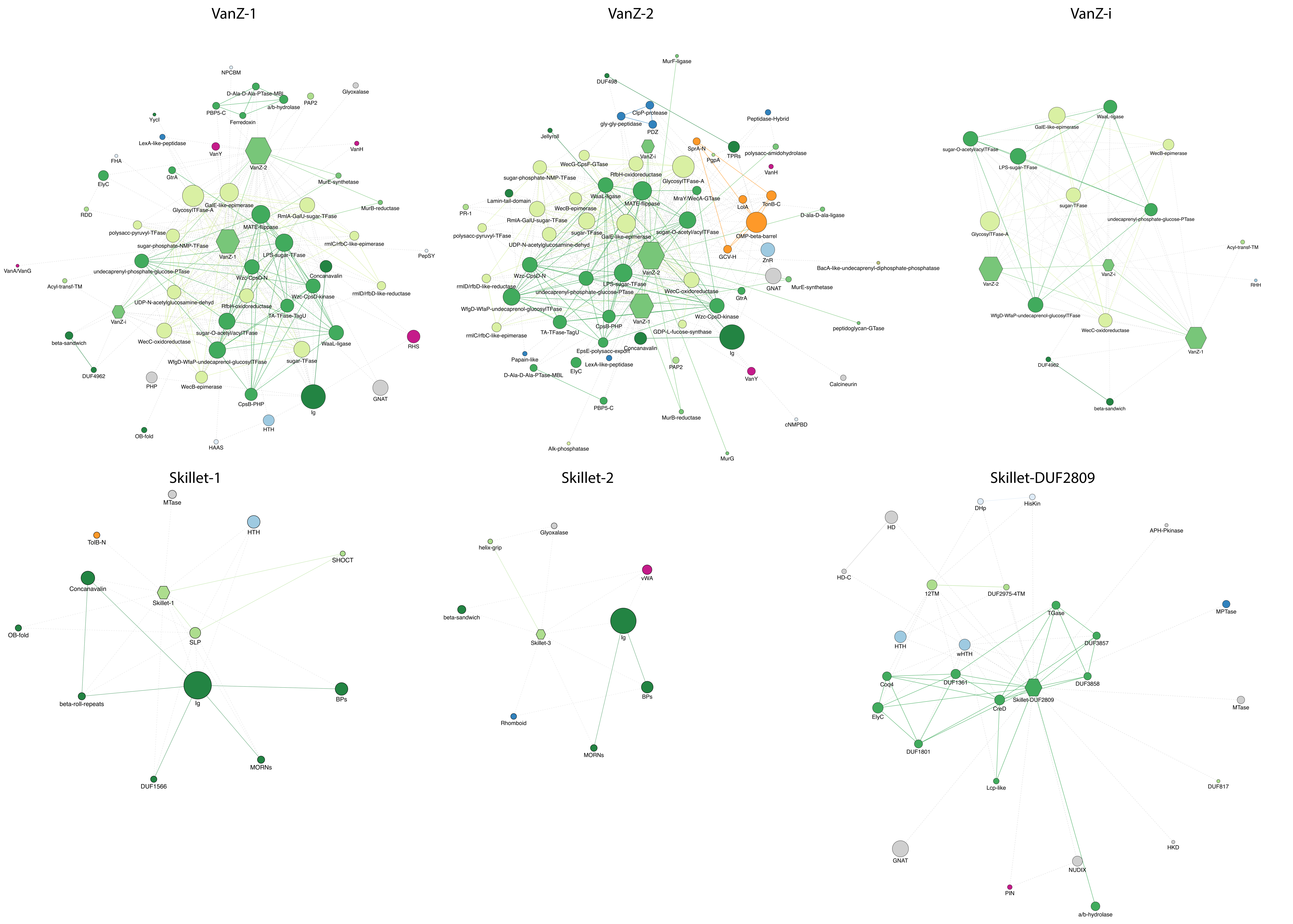

### SUPPLEMENTARY FIGURE S7.

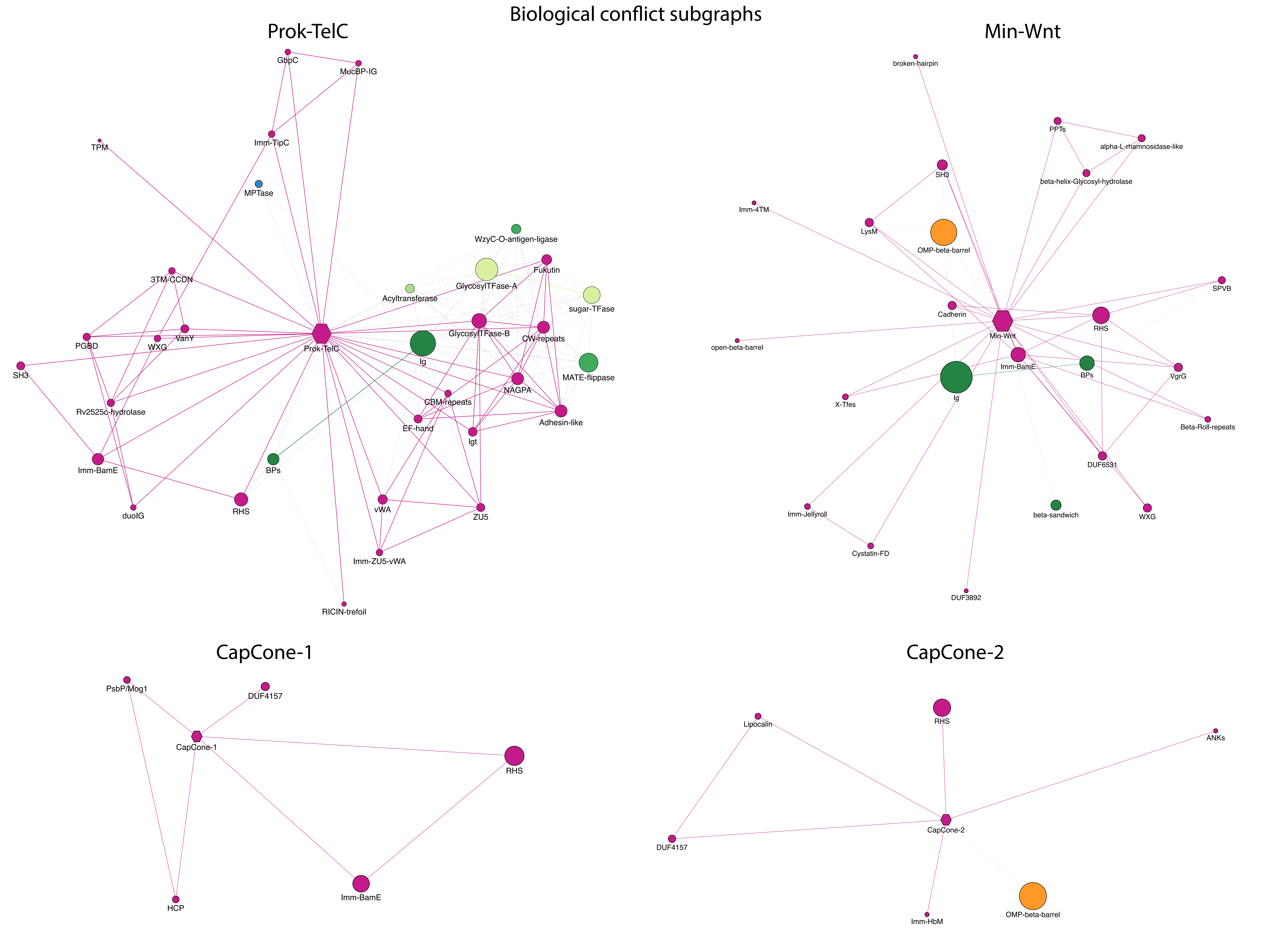

### Supplementary Figure S8. Serine-containing lipobox alignment

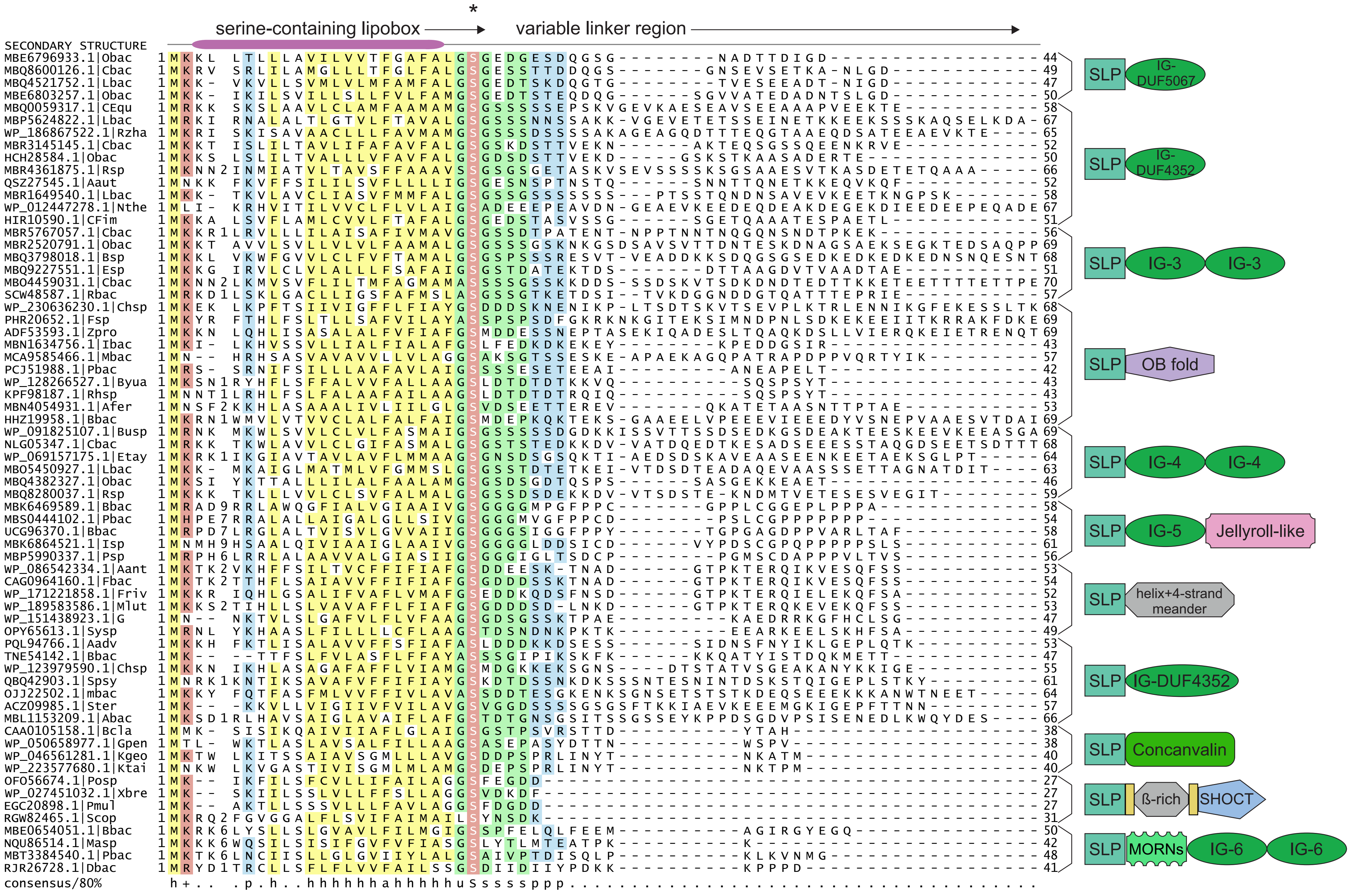

**SUPPLEMENTARY FIGURE S9.**

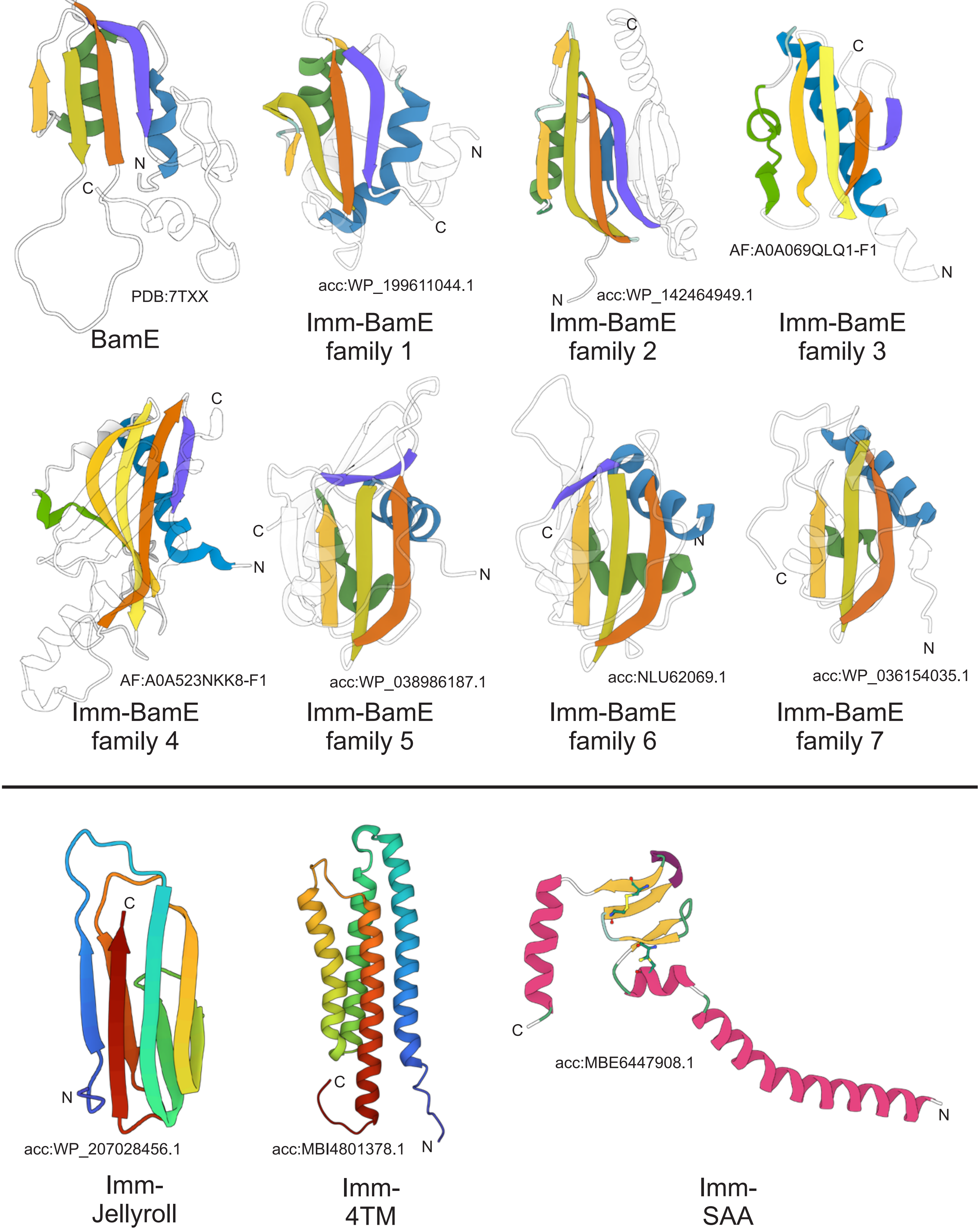

**SUPPLEMENTARY FIGURE S10.**

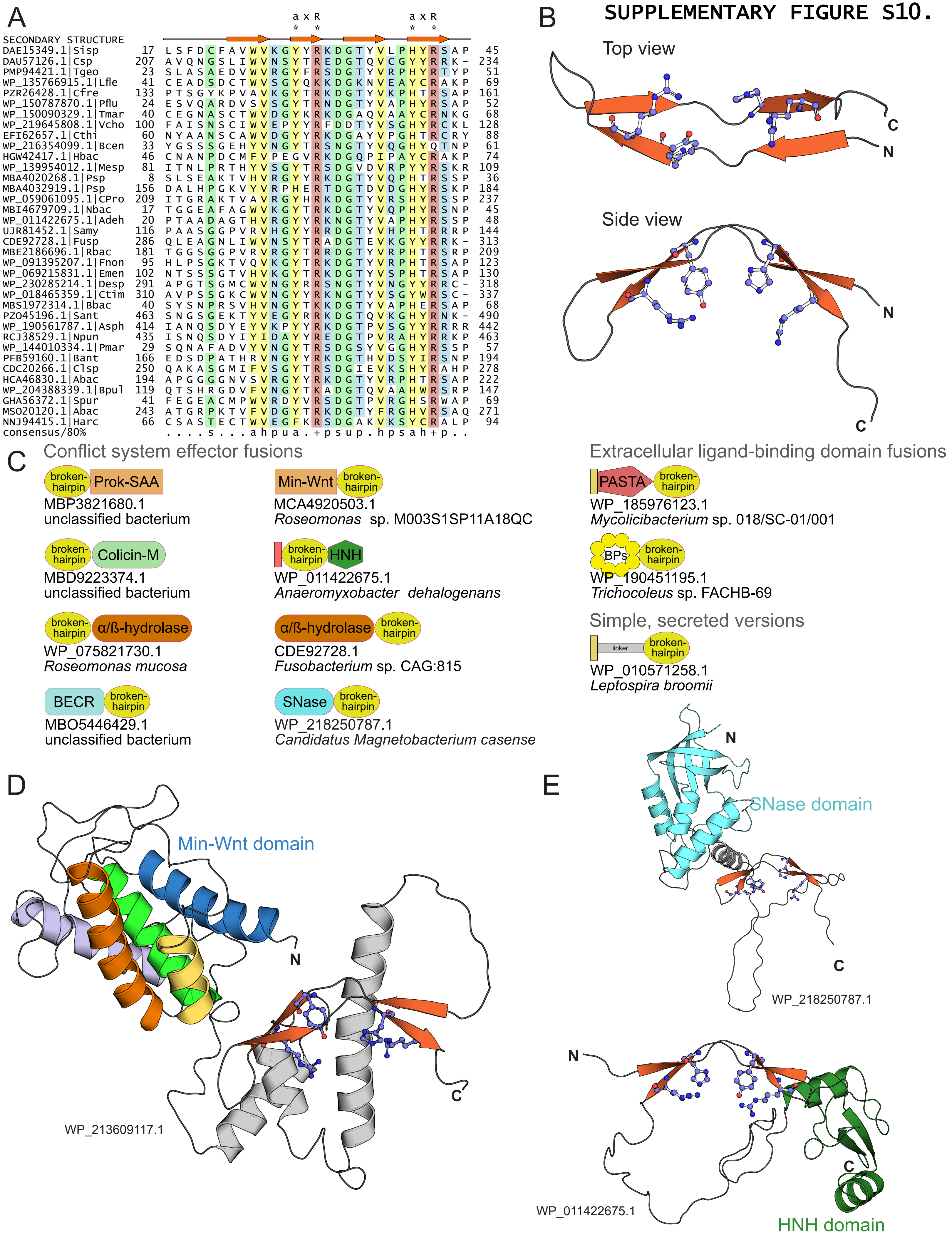
